# With our powers combined: integrating behavioral and genetic data to estimate mating success and sexual selection

**DOI:** 10.1101/129346

**Authors:** Zoé Gauthey, Cédric Tentelier, Olivier Lepais, Arturo Elosegi, Laura Royer, Stéphane Glise, Jacques Labonne

## Abstract

The analysis of sexual selection classically relies on the regression of individual phenotypes against the marginal sums of a males × females matrix of pairwise reproductive success, assessed by genetic parentage analysis. When the matrix is binarized, the marginal sums give the individual mating success. Because such analysis treats male and female mating/reproductive success independently, it ignores that the success of a male × female sexual interaction can be attributable to the phenotype of both individuals. Also, because it is based on genetic data only, it is oblivious to costly yet unproductive matings, which may be documented by behavioral observations. To solve these problems, we propose a statistical model which combines matrices of offspring numbers and behavioral observations. It models reproduction on each mating occasion of a mating season as three stochastic and interdependent pairwise processes, each potentially affected by the phenotype of both individuals and by random individual effect: encounter (Bernoulli), concomitant gamete emission (Bernoulli), and offspring production (Poisson). Applied to data from a mating experiment on brown trout, the model yielded different results from the classical regression analysis, with only a limited effect of male body size on the probability of gamete release and a negative effect of female body size on the probability of encounter and gamete release. Because the general structure of the model can be adapted to other partitioning of the reproductive process, it can be used for a variety of biological systems where behavioral and genetic data are available.

## Introduction

Sexual reproduction involves two different individuals which both invest energy in gamete encounter and possibly in offspring survival. The reproductive output of a given mating is therefore attributable to both partners. In a given population, the distribution of the reproductive success *RS*_*i,j,k*_ gained by a pair of individuals *i* and *j* on a mating occasion *k* can be summarized by a 3-dimension array of number of offspring produced between all possible pairs of males and females for each mating occasion. Then, summing such array over all the mating occasions leads to the so-called *parental table* classically used in studies of sexual selection (Arnold and Duvall 1994). An estimate of such matrix is typically generated by parentage analysis based on genetic markers (Bateman 1948, Garant et al. 2001, Avise et al. 2002, Jones and Ardren 2003, Jones et al. 2004, Serbezov et al. 2010) possibly complemented by direct observations of mating behavior (Pemberton et al. 1992, Coltman et al. 1999, Collet et al. 2014).

Classical methods in sexual selection use these parental tables to study adaptive value of traits in populations by measuring different indices of sexual selection in males and females such as opportunity for selection, selection gradients and selection differentials (Bateman 1948, Wade 1979, Wade and Arnold 1980, Crow 1989). To do so, they further reduce the matrix to its margins, individual reproductive success being the sum of offspring on the individual′s row or column, and mating success being the number of positive cells on the individual′s row or column, *i.e.* the number of different individuals with which at least one offspring was produced. Sexual selection is predicted to operate provided there is variance in reproductive success and in mating success, and a strong link between these two. Likewise, a phenotypic trait is considered to be sexually selected when it covaries with mating success.

This approach has two important caveats. First, the definition of mating as the occurrence of common offspring does not account for multiple - possibly unfertile - matings, which are part of the cost of reproduction. Second, the lack of consideration for the fundamental dependency between the mating and reproductive success of an individual and the mating and reproductive success of its mates biases the estimation of selection acting on individual traits.

An illustration of the first caveat is the wealth of definitions for individual mating success during one reproductive period (Bateman 1948, Arnold and Duvall 1994, Parker and Tang-Martinez 2005, Uller and Olsson 2008, Jones 2009, Gowaty et al. 2012, Fritzsche and Booksmythe 2013). Mating success can either be viewed as the number of copulations, the number of different individuals with which the focal individual has copulated, the number of copulations that yield progeny or the number of individuals with which progeny is produced. While the two latter definitions inform precisely on the fitness benefits, the first and second definitions also integrate potential costs, be it time, energy, predation risk, or disease transmission. Because benefits and costs are both essential to understand the evolution of sexual selection, it should be of interest to study both points of view in a single framework to estimate sexual selection indices. It is noteworthy that the definition of mating success is to a great extent constrained by methodological possibilities. Standard methodological approaches using parental tables obtained from genetic assignations can only target the fourth definition and generally produce biased estimates of it (Collet et al. 2014). These approaches deduce individual mating success by counting the number of non-zero elements on the individual line of the parental table. In this case, a zero value for a given pair can be the outcome of either pre-copulatory, post-copulatory or sampling processes: no copulation, copulation but no gamete fertilization, gamete fertilization but offspring dying before sampling, offspring alive but failing to be sampled. Similarly, a non-zero value can also carry more information than just the total reproductive success between a pair of individuals, since it can be the outcome of a variable number of matings, which is of importance to measure reproductive investment. In this perspective, matrices of copulation success as obtained by direct observations of mating behavior obviously contain data that are complementary to parentage assignation methods (Collet et al. 2014). We therefore need statistical models integrating both behavioral and genetic data to provide estimates of the various definitions of mating success, by disentangling pre-copulatory and post-copulatory components as already suggested by several authors (Arnold and Wade 1984, Pischedda and Rice 2012, Pélissié et al. 2014).

The second caveat is less evoked in the literature although intuitively simple: in sexual reproduction, reproductive success between two individuals should be attributable to both. Yet, one usually analyzes reproductive success as an individual characteristic, with no regard for the effect of the sexual partner. Classical studies only focus on the marginal sums of the parental table, and therefore cannot control for sexual partner trait or mating success variation. Selection indices are estimated by regressing the margins of the parental table against the vector of values of phenotypic traits, independently for males and females. A direct consequence is that we might detect a significant correlation between a trait and mating success or reproductive success for a sex, and interpret it as evidence of direct selection, whereas indirect selection could for instance be at work by mean of non-random association between sexual partners’ traits. We therefore need an approach in which the mating and reproductive success of a pair of individuals accounts for the phenotype of both individuals, instead of using twice the same data to draw seemingly independent conclusions.

To solve both matters, we propose a model that combines genetic data (parental table) and behavioral data (encounter and mating matrix) to 1) describe the different components of reproductive success (here encounter rate, rate of gamete release, number of offspring produced) for each mating occasion within the reproductive season, and 2) infer the joint effects of both male and female phenotype on each component of the reproductive success. The conditional structure linking the successive components of pairwise reproductive success is the key to extract information from both behavioral and genetic data: presence of offspring from a pair of parents implies encounter and gamete release, even if these are absent from behavioral data, whereas observation of gamete release despite the absence of common offspring allows distinguishing between zero-value due pre-copulatory and post-copulatory mechanisms. We illustrate the model using a reproduction experiment data for *Salmo trutta* as a case study, with body size as an example of phenotypic covariate as it is known to be involved in sexual selection in salmonids (Jacob et al. 2007, Labonne et al. 2009) and could therefore have an effect on each of these components of sexual selection. More precisely, larger males were expected to have a higher probability of encounter and mating with females because they could oust smaller males from nesting sites. In cases of multiple mating (several males ejaculate over a female’s eggs), they were also expected to sire more offspring than smaller males because their closer proximity with females during spawning gives them an advantage in sperm competition. Larger females may be expected to have a higher probability of encounter because they may attract more males than smaller females. However, larger females may not have a higher probability of mating. Because body size is highly correlated with the number of eggs, larger females were expected to produce more offspring.

## Methods

### Reproduction experiment

The experiment was conducted in semi-natural channel beside Lapitxuri stream, a tributary to the Nivelle River in south-western France (+43° 16′ 59″, −1° 28′ 54″) (De Gaudemar and Beall 1999), from November 2012 to the end of March 2013 (brown trout spawning season under this latitude). The experimental setup is the one used in the “constant environment” treatment in Gauthey et al. (2016). Three linear and communicating sections of the channel were used during the experiment, each measuring 10 meter long and 2.80 meters wide. The central section was fit out for spawning, with the appropriate gravel size (1 to 4 cm diameter), water depth (20 cm) and current speed (0.11 m.s^−1^). In the two extreme sections, a more complex environment was installed with bigger substrate size, visual obstacles (woods, bricks) and pools that provided hiding and resting areas. The parent pool consisted in 52 brown trout adults (19 males and 33 females) captured in two rivers: River Bastan (+43° 16′ 2.51″, −1° 22′ 32.46″) and River Urumea (+43° 14′ 31.81″, −1° 55′ 28.98″). Upon electrofishing, each trout was anesthetized (30 mg.l^−1^ benzocaine), sexed, measured for fork length, weighed, and photographed to allow individual identification on subsequent video recordings. On waking, fish were released in the three section of the semi-natural river, where they were free to move until the end of the experiment.

### Behavioral data

The fish were observed for at least 15 min in the morning and in the evening from the bank, in order to detect behaviors associated to spawning activity. When reproductive behaviors indicating that a female and one/or several male(s) were close to spawning (digging female, chases between males), subaquatic and aerial digital camera videos were placed in the river or on the bank in order to record the spawning act (Aymes et al. 2010, Tentelier et al. 2011).

For each observed mating occasion (one female lays her eggs and at least one male releases sperm), up to 3 hours of videos were analyzed, 1h30 before gamete release and 1h30 thereafter in order to identify individuals involved in the encounter process and in the gamete release process. To do so, a zone of one meter around the female’s nest construction was defined. Individual recognition was performed by comparing pictures took before the experiment to the image on the video. As black and red spot density and position vary consistently between individuals and do not change during the reproduction period, they were accurate tools for individual discrimination. Such discrimination was however difficult when fish were too far from the camera, in which case they were labelled as “unknown” (about 30% of observations). Individuals were considered present when they entered the zone. They were considered absent when they were outside the zone. A female and a male were considered to have encountered each other on a given mating occasion if they were both present on the zone at least once during the three-hour period. The total number of encounters observed during the experiment was stored in a males × females matrix. The simultaneous gamete release of both male and female was also stored in a males × females matrix. The term “observed mate” will be hereafter used to refer to individuals that have actually been seen copulating together. The behavioral survey ended when no reproductive behavior had been detected for one week.

### Genetic data

At emergence (800 degree.days: about two months after the last spawning event), juveniles stemming from the reproduction in the experimental channel were collected by either electrofishing or trapping at the downstream end of the experimental reach. They were anesthetized and killed under a lethal dose of 2-phenoxyethanol and placed individually in a tube of absolute ethanol (90°) upon molecular analysis. A small piece of pelvic fin was also taken on adults and stored in 90% ethanol upon molecular analysis. DNA extraction, PCR amplification and genotyping at eight microsatellite loci provided data for parentage analysis run on Cervus software (Kalinowski et al. 2007), as described in Gauthey et al. (2015). The parentage analysis resulted in the parental table, a males × females matrix figuring the number of offspring assigned to each pair.

### Classical selection analysis

Behavioral and genetic data were analyzed using classical methods. We computed the opportunity for selection, as the ratio of variance in the number of offspring genetically assigned on it squared mean. Likewise, opportunity for sexual selection was computed as the ratio of variance in the number of genetic mates on its squared mean. The term “genetic mate” is hereafter used to refer to mates deduced from genetic assignation analysis. Bateman’s gradient (*β*_*ss*_) was measured using a simple linear regression between the number of offspring assigned and number of genetic mates. To quantify selection on individual phenotype, body size was regressed against the number of encounters and the number of observed mates on videos, and on the number of offspring and number of genetic mates.

### Statistical model

The general philosophy of the model was to consider reproduction between pairs of individuals as a series of *K* mating occasions, defined as events on which at least one male x female pair mated, *i.e.* encountered, emitted gametes simultaneously and produced offspring. So, each mating occasion consisted of three successive processes: encounter (a binomial variable indicating if male *i* met female *j* on mating occasion *k*), gamete release (a binomial variable indicating if male *i* and female *j* both emitted their gametes on mating occasion *k*), and the number of offspring produced (a discrete quantitative non negative variable describing the number of offspring produced by male *i* and female *j* on mating occasion *k*). Any pair could be involved in each process of any mating occasion so the three processes could be modelled as arrays, the dimensions of which were males, females and mating occasions. The effect of male and female body size, as well as random individual effects on each process conditional of the preceding one was then assessed with Bayesian inference.

Although behavioral data stored in matrices of encounter and gamete release were only available for the *K*_*obs*_ mating occasions that were video recorded, genetic data on the number of offspring produced pool all *K* mating occasions, because offspring were sampled at the end of the spawning season, as it is often the case. Hence, a first challenge to the model was to unfold the parental table (matrix of pairwise reproductive success) *N*_*i,j*_ in *K* sub matrices, with *K* the total number of mating occasions that occurred in the mating season. We simply assumed that 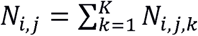. However, behavioral data are generally incomplete: here the total number of mating occasions K (*K*_*obs*_ ≤ *K*) as well as the probability *p*_*o*_ to observe encounter between a male *i* and a female *j* at each of the *K*_*obs*_ known mating occasions must be estimated. For the probability of observation, the occurrence of an observed encounter *OE*_*i,j,k*_ was modeled as *OE_i,j,k_ = E_i,j,k_ x O_i,j,k_*, where *E*_*i,j,k*_ and *O*_*i,j,k*_ were both binomial variables sampled in Bernoulli distributions of mean *p*_*e*_ and *p*_*o*_, respectively the probability that the encounter happened and the probability that it was observed. A zero *O*_*i,j,k*_ meant we had no direct behavioral data, so encounter rate and rate of gamete release could not be directly estimated. In such case, we simply simulated the expected behavioral data using the posterior densities from estimated parameters for the *K*_*obs*_ mating occasions where behavioral data were known. The total number of mating occasions, *K*, could be estimated directly in the model because the posterior distribution revealed the best combination of behavioral and genetic data conditional on the value of *K*. When behavioral data were re-simulated from their posterior distribution, the value of *K* could therefore be jointly estimated.

We tested the additive effects of male and female body size (*BS*_*i*_ and *BS*_*j*_) on encounter rate (*E*_*i,j,k*_), rate of gamete release (*G*_*i,j,k*_) and offspring number (*N*_i*,j,k*_) as following:

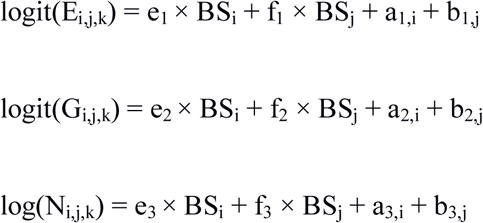

where *a*_*.,i*_ and *b*_*.,j*_ were male and female random effects, which were included to account for the fact that each individual could be involved in several mating occasions during the season. *e*_*1*_, *e*_*2*_, *e*_*3*_ are the male body size effects on encounter rate, rate of gamete release, and offspring number respectively, and *f*_*1*_, *f*_*2*_ and *f*_*3*_ are the female body size effects likewise.

Statistical inference was conducted in the Bayesian framework under JAGS 4.1.0 (Plummer 2003). Two independent MCMC samples of 10000 draws with a thinning of 100 were used, with 5000 draws as a burning period, and another 5000 draws to obtain posterior estimates. Chain convergence was checked using the Gelman-Rubin test (Gelman and Rubin 1992). In each chain, we used non informative Gaussian and independent prior distributions (mean = 0, variance = 1000) for hyperparameters: *e*_*1*_, *e*_*2*_, *e*_*3*_, *f*_*1*_, *f*_*2*_, *f*_*3*_, Beta prior distribution *B*(1,1) for *po,* Gamma distribution Γ(0.001, 0.001) for the precision of each Gaussian distribution in which random effects (*a*_1_, *a*_2_, *a*_3_, *b*_1_, *b*_2_, *b*_3_) were drawn, and a uniform distribution [15,150] for *K*. The full model code and data are available in Supplementary material Appendix 1.

## Results

### Behavioral and genetic data

Three individuals were removed from the data set because of escape from the experimental channel (2 males and 1 female). This event happened during the two first weeks of the experiment when reproductive period just started and these individuals were not observed as sexually active on the videos. These three individuals were therefore discarded from the different analyses.

In total, 22 spawning acts were video recorded (*K*_*obs*_ mating occasions) during the reproductive season. Within these *K*_*obs*_ occasions, 14 females out of 32 and 12 males out of 17 were observed, totalizing 75 pairwise encounters. Thirteen females and 7 males were observed releasing their gametes, totalizing 22 pairwise copulations (no multiple mating - where several male emit their gametes simultaneously - was observed). For five mating occasions, some individuals (1, 1, 2, 2 and 4 individuals respectively) which did not release their gametes were too far from the camera to be unambiguously identified. These individuals were therefore not taken into account for the encounter observations. Stripping at recapture showed that almost all individuals (especially females) had released their gametes by the end of the experiment (only two females did not lay their eggs), and some redds were detected in places where we did not place our cameras, indicating that a significant proportion of spawning events was not observed.

A total of 555 juveniles and 49 parents were genotyped. Among those individuals, 551 juveniles were assigned to 41 pairs of parents (10 males and 22 females) at a confidence level of 95%. Number of offspring varied from 0 to 201 in males (mean ± sd= 32 ± 64) and between 0 and 86 for females (mean ± sd= 17 ± 24). Only 12 pairs were both seen releasing gametes and assigned offspring, so joint gamete release was assessed for 29 pairs by genetic data only. At the individual level, the number of gamete releases observed on video was correlated to the number of mates inferred from the genetic analysis (Pearson’s r = 0.66, p < 0.0001). From the genetic data, the opportunity for selection was 4.49 for males and 2.34 for females. The opportunity for sexual selection was 2.69 for males and 0.81 for females. Bateman’s gradient was 17.06 for males (t = 4.229 on 15 degrees of freedom, p = 0.0008) and 13.70 for females (t = 4.175 on 30 degrees of freedom, p = 0.0002).

Using the behavioral data only, we found that male body size did not affect number of females encountered (t = 1.195 on 15 df, p = 0.251, Fig. 1a.), but it affected positively the number of mates (slope = 0.03, t = 3.268 on 15 df, p = 0.005, Fig. 1b.). Female body size affected neither the number of males encountered (t = 0.072 on 30 df, p = 0.943, Fig. 1a.) nor the number of mates (t = −0.304 on 30 df, p = 0.763, Fig. 1b.). Using the genetic data only, we found that male body size had a positive effect on number of mates (t = 3.851 on 15 df, p = 0.002, Fig. 1b.) and number of offspring (t = 0.2604 on 15 df, p = 0.003, Fig. 1c.), whereas female body size affected neither the number of mates (t = 0.659 on 30 df, p = 0.515, Fig. 1b.) nor the number of offspring (t = 0.1782 on 30 df, p = 0.845, Fig. 1c.).

**Figure 1.**
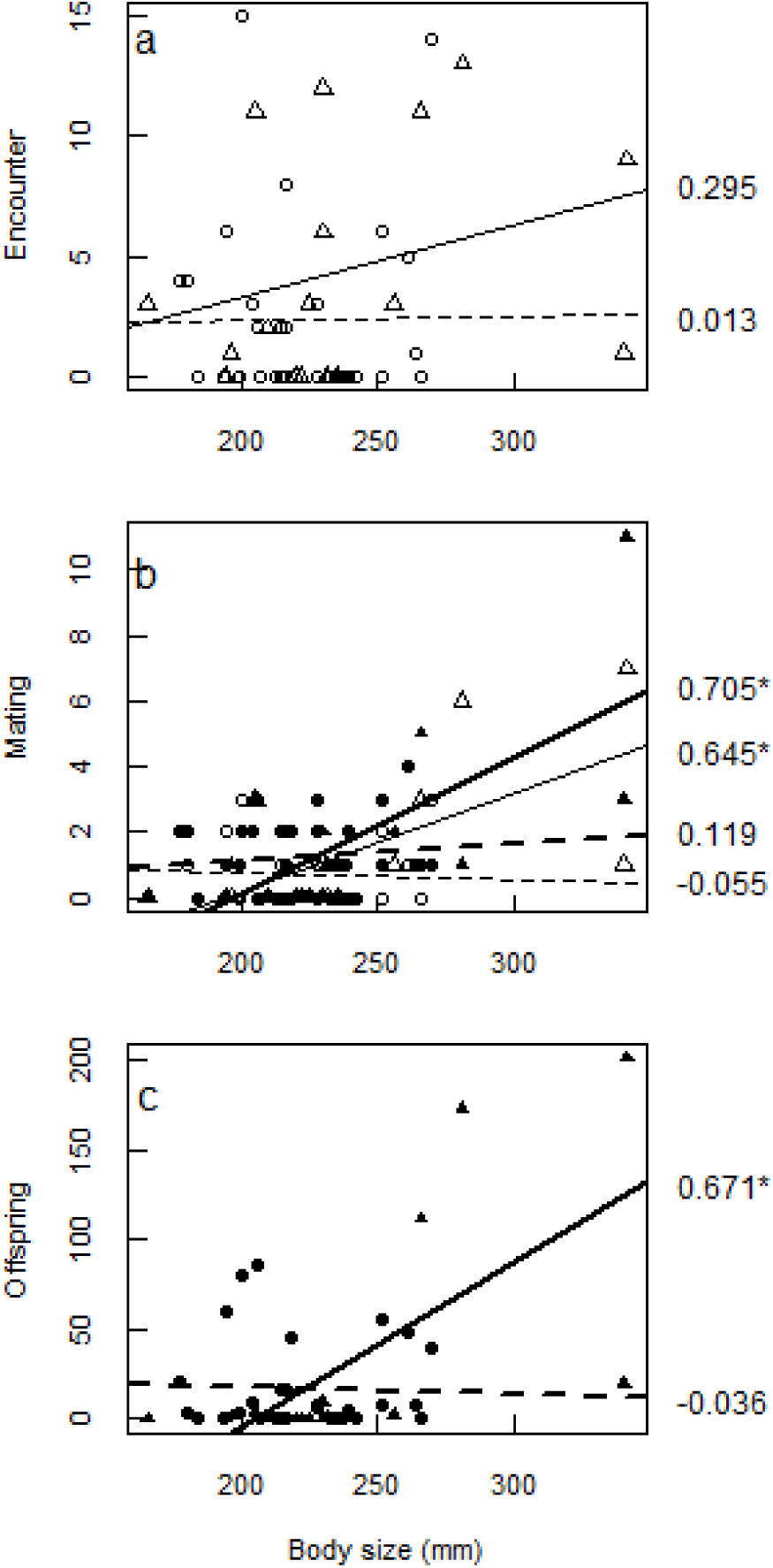
Linear regressions of brown trout body size against a) the number of individuals of the opposite sex which were encountered, b) the number of mates and c) the number of offspring assigned. Circles and dashed lines are for females, and triangles and solid lines are for males. Empty and filled symbols correspond to behavioural and genetic data, respectively. For b) mating success was measured as the number of individuals of the opposite sex with which the focal individual was observed emitting gametes (empty symbols) and as the number of individuals with which it shared offspring (filled symbol). Values on the right margin indicate the Pearson’s correlation coefficient of the corresponding regression line, with asterisk indicating p < 0.05.

### Model output

The posterior of all parameters for the model are provided in Supplementary material Appendix 2. Although only 22 pairwise gamete releases were recorded on video and 41 families were detected by genetic analysis, the posterior distribution of *K*, the number of mating occasions, had a median of 117 [1^st^ quartile = 103; 3^rd^ quartile = 132]. The posterior distribution of the probability of observing an encounter between two individuals in a given mating occasion, *p*_*o*_, had a median of 0.66 [0.63; 0.76]. Based on the joint posterior probabilities of all parameters (effects of male and female body size, and individual random effects), the model predicted an average (± standard deviation) of 47 (± 25) encounters per male, 25 (± 27) encounters per female, 9.8 (± 8.4) gamete releases per male, 5.2 (± 8.2) gamete releases per female, 32 (± 36) offspring per male and 17 (± 33) offspring per female.

Male body size had no effect on the probability of encounter or on the number of offspring produced at each mating occasion, and had a very slight positive effect on the probability of gamete release (Fig. 2). The posterior distribution of the parameter associated to the effect of male body size on gamete release (*e*_2_) had a median of 7.7428. 10^−3^, which corresponds to an odd of encounter multiplied by only 1.007 for each additional millimeter. Given that male body size ranged from 165 to 342 mm, this would predict, other things equal, a 3.7 odds ratio between the longest male and the shortest one. Female body size had a negative effect on both the probability of encounter and the probability of gamete release but did not affect the number of offspring produced (Fig. 2). The median of the posterior distributions on *f*_1_ and *f*_2_ were −0.02386 and −0.02126, resulting in odds of encounter and gamete release being multiplied by 0.976 and 0.979, respectively, for each millimeter. Given that female body size ranged from 177 to 270 mm, the odds ratio between the longest and the shortest female would be 0.11 for encounter and 0.14 for gamete release.

**Figure 2.**
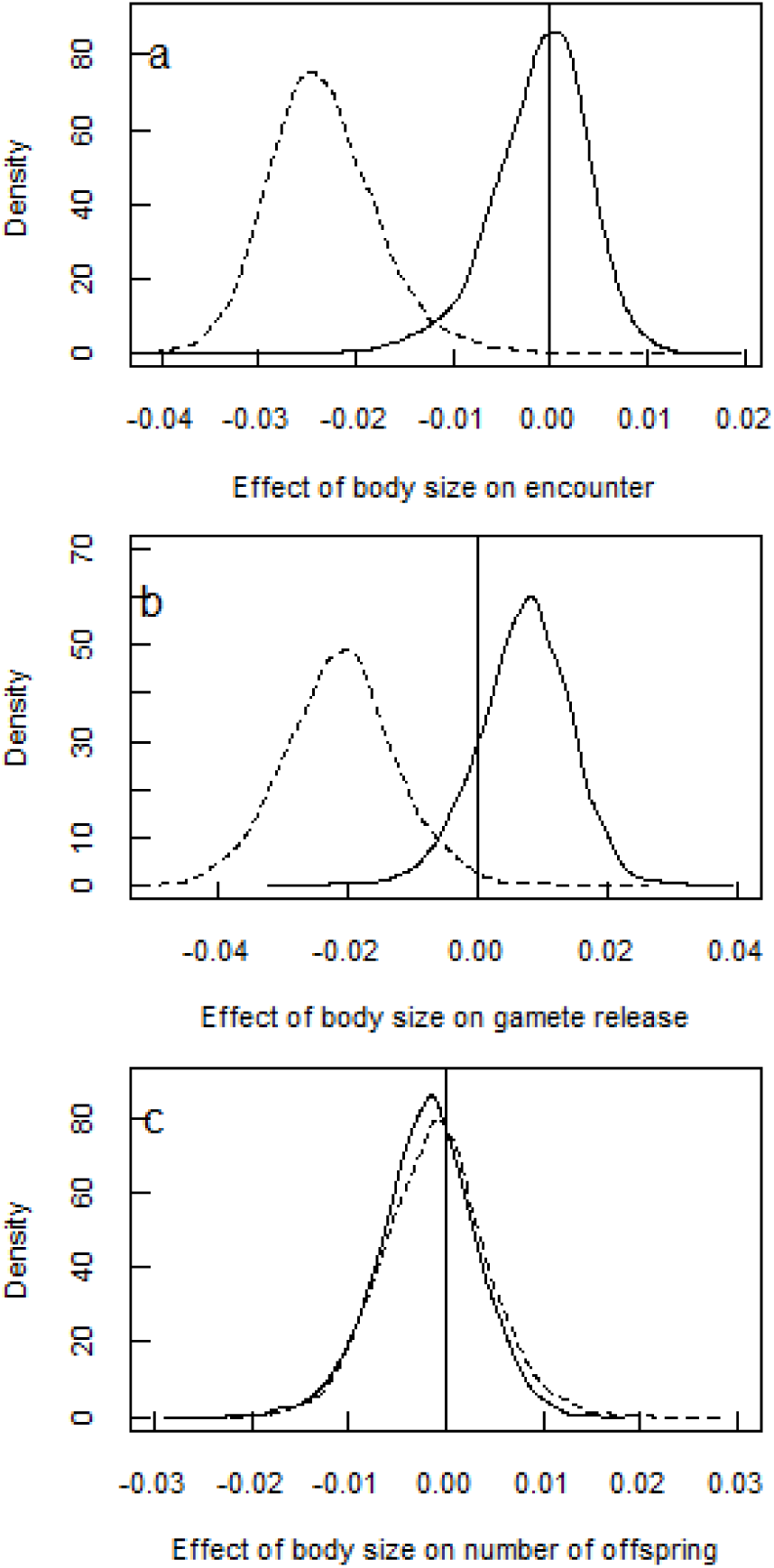
Posterior probability distributions of model parameters associated to the effect of brown trout body size on a) the probability of encounter, b) the probability of gamete release and c) the number of offspring produced on each mating occasion. Dashed and solid lines are for females and males, respectively

Random effects were more variable for females than for males for the probability of encounter and the probability of gamete release, while male random effects were more variable than female’s for the number of offspring (Fig. 3). Moreover, random effects on probability of encounter, probability of gamete release and number of offspring were positively correlated for both sexes (Fig. 3). Because random effects for the probability of encounter and gamete release act on the logit scale and random effects for the number of offspring act on its logarithm, they should be interpreted such that individuals having a random effect of 0.5, 1, 2 or 4 have 1.6, 2.7, 7.4 or 54.6 times higher odds or more offspring than the average individual, respectively.

**Figure 3.**
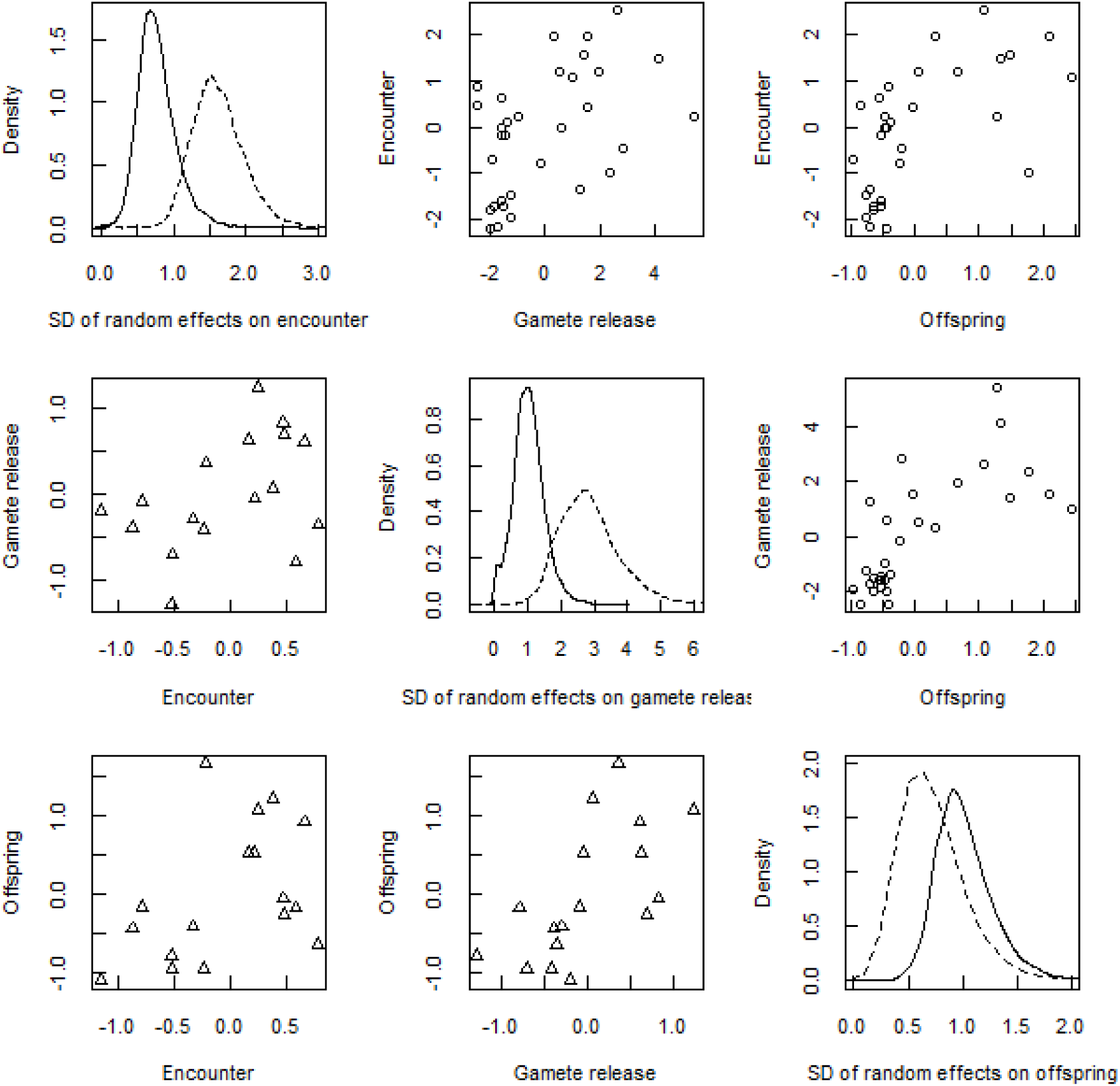
Random individual effects on the probability of encounter, the probability of gamete release and the number of offspring produced by brown trout on each mating occasion. The diagonal indicates the posterior probability distribution of the standard deviation of the Gaussian distribution in which random effects for the three components of reproductive success were drawn (dashed and solid lines are for females and males, respectively). Plots above the diagonal show the pairwise relations between random individual effects on each process, for females (one circle per female). Plots below the diagonal show the same thing for males (one triangle per male).

Joint posterior probability distributions were used to predict the number of encounters, gamete releases and offspring for each individual and these predictions were plotted against the number of encounters and gamete releases observed on videos and number of offspring genetically assigned (Fig. 4). In most cases, numbers predicted by the model exceeded the number of observations, but the number of offspring predicted by the model could be smaller than the number of offspring actually assigned, especially for females.

**Figure 4.**
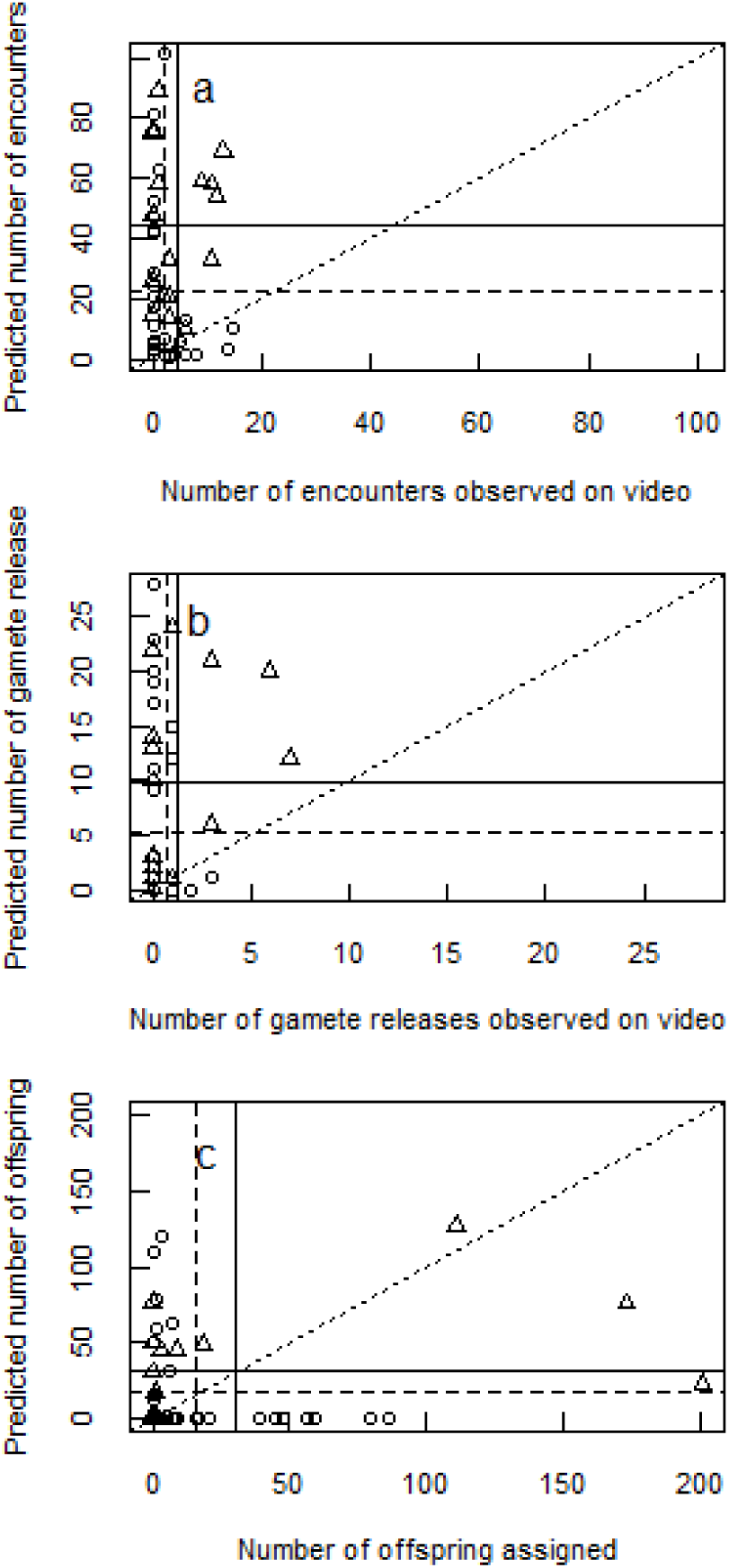
Predictions based on the joint posterior distributions of model parameters, against values observed in the raw data for a) the number of encounters, b) the number of gamete releases and the number of offspring assigned to individual brown trout. Circles and triangles are for females and males respectively. Dashed and solid lines indicate, for females and males respectively, the mean of each variable. The dotted line has intercept zero and slope one, which would correspond to a perfect fit between observed and predicted values.

## Discussion

In this study, we used two approaches to estimate the effect of a phenotypic trait (here body size as an example in brown trout) on different components of sexual selection. Both approaches lay on behavioral observation of encounter and mating, and genetic assignation of offspring. On the one hand, we applied classical analyses on data pulled out from the marginal sums of each male × female matrix: number of encounters and gamete releases observed on videos, and number of offspring and mates inferred from genetic assignation. There we found that body size, in males only, would correlate positively with mating success and offspring number, but not with encounter rate. On the other hand, we developed a statistical framework combining all these data, thereby enabling information to circulate through the successive processes of encounter, gamete release and offspring production. This new approach accounted for the three-dimensional structure of the data: males, females and mating occasions. This allowed a qualified definition of mating success and disentangling the joint effects of male and female phenotypes on the different components of reproductive success. There we found that body size, in females only, would correlate negatively with encounter rate and mating success, but not with offspring number.

### What is mating success?

The multiple definitions of mating success have been shaped by a dichotomy of approaches, which our model aimed at overcoming. On the one hand, because the classical approach based on the genetic parental table is oblivious to both ineffective mating acts and multiple inseminations between the same pair of individuals, it has constrained the definition of mating success to the number of individuals with which the focal individual produces offspring that are alive at sampling (Arnold and Duvall 1994). On the other hand, the not less classical approach based on the sole observation of copulatory behavior, unable to access the reproductive output, focused the definition of mating success on the number of copulations or number of copulatory partners. By combining behavioral and genetic data in a common framework, our analysis embraced multiple aspects of mating success. The combination of genetic data and behavioral observations to account for mating acts the offspring of which were not sampled was also adopted for instance by Collet et al. (2014) and Pélissié et al. (2014) but their approach relied on complete knowledge of copulation events in the mating group to disentangle the contribution of pre-copulatory and post-copulatory components of reproductive success. Our approach consisted in merging the behavioral and genetic datasets, both incomplete – a common situation in ecology and evolution -, and took advantage of the conditional structure of the successive components of mating success: encounter, simultaneous gamete release and offspring production.

At the scale of the reproductive group, our behavioral observations showed 75 male × female encounters and 22 pairwise gamete releases, whereas the parental table based on genetic assignation indicated that 41 broods were produced. Given that only 12 pairs both were observed copulating and had their offspring sampled, a rough estimate of the probability that a pair was observed mating would be 12/41 = 0.29, and a rough estimate of the probability of a pair having its offspring sampled would be 12/22 = 0.54. This would mean that 12/(0.29*0.54) = 76 matings had occurred, 10 of which were video recorded only, 29 of which were detected genetically only, 12 of which were detected both on video and by the genetic analysis, and 25 were missed by both methods. In our model, the parameter *K*, called the number of mating occasions, was estimated to be 117, meaning that each pair had 117 occasions to mate. This concept of mating occasion, defined as an event on which any male x female pair *may* encounter, emit gametes and produce offspring, was much broader than mating, defined as an event on which a male x female pair *does* encounter, emit gamete and produce offspring. By splitting individual mating success in a number of mating occasions (trials), our modelling approach considered mating success as the result of a Bernoulli process, with inferences made on the probability of success. Moreover, this success of joint gamete release was conditioned on the success of encounter on each occasion, and conditioned in turn the number of offspring produced. This conditional structure is in line with the concepts of “sexual networks” and “sexual niche” (McDonald et al. 2013, Ziv et al. 2016), which acknowledge that an individual interacts with (competes with, courts, chooses among) only a subset of the population. Hence, sexual selection should be measured among individuals that actually interact.

Individual variance in mating success is the fuel for sexual selection, and the opportunity for sexual selection is computed as the variance in number of mates on the squared mean of number of mates. Based on classical treatment of genetic data, opportunity for sexual selection was higher for males (2.69) than for females (0.81) as usually expected (Bateman 1948). However, our model indicated that both the probability of encounter and the probability of gamete release on a mating occasion was more variable among females than among males, since the effect of body size (Fig. 2) and the individual random effects (Fig. 3) on theses probabilities were larger for females than for males. This counter-intuitive result may be due to the model detecting a higher mates number (gamete releases) than the sole genetic approach. Moreover, for both sexes random effects on the probability of encounter, the probability of gamete release and the number of offspring produced were positively correlated. This suggests that some individuals performed consistently better than others for the three processes, *i.e.* had a higher probability of encounter, a higher probability of gamete release once a partner was encountered, and a higher number of offspring produced once mated, unconditional on body size.

### Combined effects of male and female phenotype on the components of reproductive success

Sexual selection on phenotypic traits is classically quantified for each sex separately, by regressing the number of mates against phenotypic trait in a separate model for each sex (Andersson 1994). Here, the statistical unit is the individual, and individual mating success and reproductive success are assumed independent among individuals. However, mating and reproduction are essentially matters of pair, hence both male and female traits contribute to pairwise mating success and reproductive success on a given occasion. Our approach was therefore to consider the mating occasion as the statistical unit, and infer the effect of traits (here, body size) borne by individuals involved in that occasion on its outcome. This approach departs from selection theory, to which regression models fit well (Price 1970, Lande and Arnold 1983, Moorad and Wade 2013), but allows insight on the mechanisms by which traits affect reproductive success.

Applying classical linear regressions to our data indicated that larger males tended to have more encounters, and had significantly more gamete releases, more genetic mates and more offspring, while female body size affected none of the behavioral or genetic indicators of reproductive success. However, our model accounting for the size of both males and females as well as individual random effects on each reproductive process indicated that larger females had a lower probability of encounter with males and a lower probability of gamete release, whereas male size affected neither encounter, gamete release nor number of offspring. Hence the output of the two analyses differed a lot.

The difference between the linear regression approach and ours is due to three features of our model which lack in the classical approach: 1) conditioning of each process (encounter, gamete release and offspring production) on the preceding one, 2) simultaneous estimation of the effect of male and female phenotype, and 3) random individual effects. The conditional structure of the model allowed to infer the effect of individual phenotype on each process independently, whereas regression made on all individuals may confound them. For instance, our analysis indicated that larger males tended to have a higher probability of releasing gametes with the females they encountered, but once mated they did not tend to sire more offspring. According to the regression analysis, though, the number of offspring was positively related to body size, but this relationship was indirect and mediated by the positive relation between number of mates and number of offspring (Bateman gradient). Although reproductive success may be split into multiplicative components on each of which individual phenotype can be regressed, such analysis requires as many regressions as components (e.g. Arnold and Wade 1984, Tentelier et al. 2016), whereas our model encompasses them all.

By considering the mating occasion as the statistical unit, we assumed that the realization of each process was potentially attributable to both sexual partners, thereby decomposing the variance between both male and female body size effects. The consequences on the results are rather strong, since for instance, we detected that female body size was then negatively impacting both encounter and mating processes. Additionally, the use of random effects on process further avoided to falsely attribute variance to body size. The classical approach – which implies a pseudo-replication effect since the data are used twice, once for males, once for females - could see no effect of female body size.

Now as to why female body size, for instance, had a negative effect on encounter and gamete release probability, and no positive effect on offspring number, we must turn to the behavioral knowledge of the species. In particular, assortative or disassortative encounter and mating, be it the result of mate choice, intrasexual competition or chance, is possible in brown trout (Petersson et al. 1999, Labonne et al. 2009): bigger females tend to be aggressively monopolized by bigger males, thereby limiting their access to a higher number of potential mates. Unfortunately our dataset is too small to properly infer the effect of interaction between male and female phenotype on the different components of reproductive success (Moshgani and Dooren 2011), though it is very easy to implement in the model.

### Further applications of the model

The experimental design and the quantity of data we used to illustrate our model indubitably constrained the analysis we carried out, and one can wonder how the model can be transposed to other systems, with other types of data on either the components of reproductive success or traits affecting them. For instance, because we sampled all offspring at the end of the experiment, the genetic data did not inform much on the number of offspring produced at each mating occasion. However, in other systems where clutches are well separated in time or space, even within a reproductive season, the parental table of genetic data would also be three-dimensional (male × female × occasion) and inferences on each component of reproductive success would probably be more accurate. Also, depending on the system studied, reproductive success may be further decomposed, and inference might be done on individual or environmental features affecting the additional components. For example, one may disentangle copulation from gamete fertilization by combining behavioral data and single-molecule PCR and genotyping of zygotes just after copulation. Here, an additional three-dimension matrix containing gamete fertilization of each male-female pair at each occasion would be built, and fertilization success would be included in the model, conditioned by copulation success, and conditioning the number of offspring. This would disentangle fertilization success from zygote survival, something we were not able to do in our case study on brown trout.

Regarding traits affecting components of reproductive success, we illustrated our approach with body size only, a trait which is known to affect intrasexual competition and intersexual preference in brown trout and other salmonids (Foote and Larkin 1988, Foote 1989, de Gaudemar 1998, Blanchfield and Ridgway 1999, Fleming and Reynolds 2004, Labonne et al. 2009). Other traits could have been used, like color, which is known to play a role in brown trout reproductive success (Jacquin et al. submitted, Wedekind et al. 2008). In particular, dynamic traits could be included in our framework, since the statistical unit in our analysis is the mating occasion. Indeed, an individual could be allowed to bear a different trait value on each mating occasion, such as mating experience (Saleem et al. 2014), the outcome of previous intrasexual contests (Hsu et al. 2006), energy stores (Gauthey et al. 2015). For example, sperm depletion may lead to reduced number of offspring sired by a male on late mating occasions without affecting probability of copulation (Damiens and Boivin 2006). Finally, each mating occasion may be characterized by a given environment which could affect each component of reproductive success, either directly or in interaction with individual phenotype. For instance, water turbidity may relax sexual selection on fish coloration (Seehausen et al. 1997, Candolin et al. 2007). Likewise, individual location and wind or water current on each day of the reproductive season may have an interactive effect on pairwise reproductive success through the probability of encounter between gametes (Dow and Ashley 1998, Kregting et al. 2014).

Beyond the analysis of experimental data, the parameters estimated in a model such as the one presented here can readily be included in individual based models of sexual interaction, which implement mating as a stochastic process the success of which may be influenced by the phenotype of both individuals involved (Piou and Prévost 2012, Courtiol et al. 2016). Hence, we hope our approach will facilitate the interaction between experimental and theoretical work on sexual selection.

## Acknowledgements

This study was funded by the INTERREG Atlantic Aquatic Resource Conservation program (AARC) funded by the European Union

## Reference to supplementary material

Supplementary material (Appendix oik.XXXXX at www.oikosjournal.org/readers/appendix). Appendix 1-2.

### Supplementary material Appendix 1

JAGS code (including data) for the model inferring the effect of brown trout body size on consecutive processes of reproductive success.

Gauthey, Z. et al. 2017. With our powers combined: integrating behavioral and genetic data to estimate mating success and sexual selection. – Oikos 000: 000-000

~~~
## Supplementary File 1: model code and data.
## Gauthey, Z. et al. 2017. With our powers combined: integrating behavioral and genetic data to estimate mating success and sexual selection. Oikos 000: 000-000
#### JAGGS 4.0 CODE
~~~

~~~
model {
# likelihood
~~~

~~~
#### ENCOUNTER PROCESS
~~~

~~~
for (i in 1:I) {
     for (j in 1:J) {
     # inference of male and female body size on encounter probability
     # includes random effects
     logit(pe[i,j])<- e[1]*TM[i]+f[1]*TF[j]+a[1,i] + b[1,j]
     for (k in 1:Kobs) {
          # actual meeting process, pe=encounter probability
          E[i,j,k] ∼dbern(pe[i,j])
          # noise process, po= detection probability
          O[i,j,k]∼dbern(po)
       }
     }
}
~~~

~~~
# data fit for observed encounters
for (i in 1:I) {
     for (j in 1:J) {
     for (k in 1:Kobs) {
          # observed encounters are products of actual meeting and detection
          OEinter[i,j,k]<-O[i,j,k]*E[i,j,k]
     }
     # decomposing the observed encounter matrix
     OES[i,j]<-sum(OEinter[i,j,])
     OE[i,j]∼dnorm(OES[i,j],100)
   }
}
~~~

~~~
# data generation for non-observed encounters
for (k in (Kobs+1):Kmax) {
     for (j in 1:J) {
          for (i in 1:I) {
               E[i,j,k] ∼ dbern(pe[i,j])
          }
     }
}
~~~

~~~
#### GAMETE RELEASE PROCESS
~~~

~~~
# observed gamete releases
for (i in 1:I) {
     for (j in 1:J) {
          for (k in 1:Kobs) {
               # can release gametes only if encounter happened
               G[i,j,k]<-E[i,j,k]*GE[i,j,k]
               # probability to release gametes
               GE[i,j,k]∼dbern(pg[i,j,k])
               # inference of male and female body size on gamete release probability
               # includes random effects
               logit(pg[i,j,k])<-
e[2]*TM[i]+f[2]*TF[j]+a[2,i] + b[2,j]
                    }
               # decomposing the observed mating matrix
               Gsum[i,j]<-sum(G[i,j,])
               Gcumul[i,j]∼dnorm(Gsum[i,j],100)
               }
}
~~~

~~~
#non-observed releases
for (i in 1:I) {
     for (j in 1:J) {
          for (k in (Kobs+1):Kmax) {
               # can release gametes only if encounter happened
               G[i,j,k]<-E[i,j,k]*GE[i,j,k]
               # probability to release gametes
               GE[i,j,k]∼dbern(pg[i,j,k])
               # inference of male and female body size on gamete release probability
               # includes random effects
~~~

~~~
               logit(pg[i,j,k])<-
e[2]*TM[i]+f[2]*TF[j]+a[2,i] + b[2,j]
                              }
               }
           }
~~~

~~~
           #### OFFSPRING NUMBER PROCESS
~~~

~~~
           # offspring number for observed gamete releases
           for (i in 1:I) {
                for (j in 1:J) {
                     for (k in 1:Kobs) {
                          # can release gametes only if encounter AND gamete release happened
                          SRreal[i,j,k]<-
E[i,j,k]*G[i,j,k]*Nreal[i,j,k]
                          # Number of offspring produced
                          Nreal[i,j,k]∼dpois(pn[i,j,k])
                          # inference of male and female body size on number of offspring produced
                          # includes random effects
                          log(pn[i,j,k])<-
e[3]*TM[i]+f[3]*TF[j]+a[3,i] + b[3,j]
~~~

~~~
                          }
                }
  }
  # offspring number for non-observed gamete releases
  for (i in 1:I) {
       for (j in 1:J) {
            for (k in (Kobs+1):Kmax) {
                 # remove data that are generated above the estimate of actual total mating occasions number (RN)
                 counter[i,j,k]<-step(RN-k)
                 # can release gametes only if encounter AND gamete release happened
                 SRreal[i,j,k]<-        counter[i,j,k]* E[i,j,k]*G[i,j,k]*Nreal[i,j,k]
                 # Number of offspring produced
                 Nreal[i,j,k]∼dpois(pn[i,j,k])
                 # inference of male and female body size on number of offspring produced
                 # includes random effects
                 log(pn[i,j,k])<-
e[3]*TM[i]+f[3]*TF[j]+a[3,i] + b[3,j]
                              }
                }
     }
     # decomposing the parental table for offspring number
     for (i in 1:I) {
          for (j in 1:J) {
               SR[i,j]∼dnorm(mu[i,j],1000)
               mu[i,j]<-sum(SRreal[i,j,1:Kmax])
          }
     }
     # estimating the real number of mating occasions (somewhere between Kobs and Kmax)
     RN∼dpois(MRN)
~~~

~~~
     # Calculating male margins for encounter matrix,mating matrix, and offspring number parental table
     for (i in 1:I) {
          Emale[i]<-sum(E[i,])
          Gmale[i]<-sum(G[i,])
          Rmale[i]<-sum(SRreal[i,])
~~~

~~~
   }
~~~

~~~
   # Calculating female margins for encounter matrix, mating matrix, and offspring number parental table
   for (j in 1:J) {
        Efemale[j]<-sum(E[,j,])
        Gfemale[j]<-sum(G[,j,])
        Rfemale[j]<-sum(SRreal[,j,])
   }
~~~

~~~
   # random effects for males on encounter,gamete release, and offspring number production processes.
~~~

~~~
   for(i in 1:I) {
        a[1,i]∼dnorm(0,taum1)
        a[2,i]∼dnorm(0,taum2)
        a[3,i]∼dnorm(0,taum3)
   }
~~~

~~~
   # random effects for females on encounter,gamete release, and offspring number production processes.
~~~

~~~
   for(i in 1:J) {
       b[1,i]∼dnorm(0,tauf1)
       b[2,i]∼dnorm(0,tauf2)
       b[3,i]∼dnorm(0,tauf3)
   }
~~~

~~~
# priors
     # we know from independent data that detection probability is not close from 0 nor from 1
     # so we use an informative prior
     po∼dbeta(50,30)
     # we know from literature that middle size brown trout do not spawn a large number of time on average
     # we also know       that at least 22 mating occasions were observed.
     # so we use an informative prior
     MRN∼dunif(23,150)
     # non-informative prior for male and female body size effects on encounter,gamete release, and offspring number production processes.
~~~

~~~
     for(i in 1:3) {
          e[i]∼dnorm(0,0.001)
          f[i]∼dnorm(0,0.001)
     }
~~~

~~~
     # non-informative prior for male and female random effects on encounter,gamete release, and offspring number production processes.
     taum1∼dgamma(0.001,0.001)
     taum2∼dgamma(0.001,0.001)
     taum3∼dgamma(0.001,0.001)
     tauf1∼dgamma(0.001,0.001)
     tauf2∼dgamma(0.001,0.001)
     tauf3∼dgamma(0.001,0.001)
}
~~~

~~~
#### DATA
~~~

~~~
# female size
TF<-
c(228,218,214,232,236,199,238,216,237,204,228,242,252,266,216, 215,261,184,207,252,193,239,264,214,235,194,270,206,177,180,20 0,212)
# male size
TM<-
c(230,225,342,230,341,235,165,194,196,266,209,205,281,222,231, 220,253)
# estimated offpsring number from parental table
# males are in rows and females in columns, ordered as in body size vectors
SR<- structure(c(0,0,14,0,17,0,0,0,0,0,0,0,0,0,0,0,0,0,0,0,0,0,0,0, 0,0,0,0,0,0,0,0,
0,0,0,0,0,0,0,0,0,0,0,0,0,0,0,0,0,0,0,0,0,0,0,0,0,0,0,0,0,0,0,0,
2,0,1,0,0,0,2,0,0,8,0,0,7,1,0,0,33,0,0,27,0,0,0,0,0,0,39,0,0,2,79,0,
0,0,0,0,0,0,0,0,0,0,0,0,0,0,0,0,0,0,0,0,0,0,0,0,0,0,0,6,3,0,0,0,
3,0,0,0,0,0,0,0,0,0,7,0,0,0,0,0,9,0,0,0,0,0,0,0,0,0,0,0,0,0,0,0,
0,0,0,0,0,0,0,0,0,0,0,0,0,0,0,0,0,0,0,0,0,0,0,0,0,0,0,0,0,0,0,0,
0,0,0,0,0,0,0,0,0,0,0,0,0,0,0,0,0,0,0,0,0,0,0,0,0,0,0,0,0,0,0,0,
0,0,0,0,0,0,0,0,0,0,0,0,0,0,0,0,0,0,0,0,0,0,0,0,0,0,0,0,0,0,0,0,
0,0,0,0,0,0,0,0,0,0,1,0,0,0,0,0,0,0,0,0,0,0,0,0,0,0,0,0,0,0,0,0,
0,0,0,0,1,0,0,0,0,0,0,0,0,0,0,0,0,0,0,0,0,4,7,0,0,59,0,40,0,0,0,0,
0,0,0,0,0,0,0,0,0,0,0,0,0,0,0,0,0,0,0,0,0,0,0,0,0,0,0,0,0,0,0,0,
0,0,0,0,0,0,0,0,0,1,0,0,0,0,0,0,0,0,0,1,0,0,0,0,0,0,0,0,0,1,0,0,
0,44,15,0,0,3,0,0,0,0,0,0,0,0,17,0,5,0,0,28,0,1,0,0,1,0,0,40,18,0,1,0,
0,0,0,0,0,0,0,0,0,0,0,0,0,0,0,0,0,0,0,0,0,0,0,0,0,0,0,0,0,0,0,0,
1,0,0,0,0,0,0,0,0,0,0,0,0,0,0,0,0,0,0,0,0,0,0,0,0,0,0,0,0,0,0,0,
0,0,0,0,0,0,0,0,0,0,0,0,0,0,0,0,0,0,0,0,0,0,0,0,0,0,0,0,0,0,0,0,
0,1,0,0,0,0,0,0,0,0,0,0,0,0,0,0,1,0,0,0,0,0,0,0,0,0,0,0,0,0,0,0),.Dim=c(17L,32L))
# observed encounters in the experiment. Matrix structured as for offspring numbers
OE<-
structure(c(0,0,1,0,0,0,0,0,0,0,0,0,0,0,2,0,0,0,0,1,0,0,0,0,0, 2,0,0,0,0,0,0,
0,0,0,0,0,0,0,0,0,1,0,0,0,0,0,0,0,0,0,0,0,0,0,0,0,0,1,0,0,1,0,0,
0,0,0,0,0,0,0,0,0,1,0,0,0,0,1,0,0,0,0,1,0,0,0,0,0,0,3,0,0,0,3,0,
0,0,0,0,0,0,0,0,0,0,1,0,0,0,1,0,1,0,0,0,0,0,0,0,0,1,3,0,1,1,3,0,
0,0,0,0,0,0,0,1,0,0,0,0,0,0,0,0,0,0,0,0,0,0,0,0,0,0,0,0,0,0,0,0,
0,0,0,0,0,0,0,0,0,0,0,0,0,0,0,0,0,0,0,0,0,0,0,0,0,0,0,0,0,0,0,0,
0,0,1,0,0,0,0,0,0,0,0,0,0,0,0,0,1,0,0,0,0,0,0,0,0,0,0,0,0,0,1,0,
0,0,0,0,0,0,0,0,0,0,0,0,0,0,0,0,0,0,0,0,0,0,0,0,0,0,0,0,0,0,0,0,
0,0,0,0,0,0,0,0,0,0,0,0,0,0,0,0,0,0,0,0,0,0,0,0,0,0,0,1,0,0,0,0,
0,0,0,0,0,0,0,1,0,0,0,0,0,0,1,0,1,0,0,0,0,0,1,0,0,2,2,0,0,0,3,0,
0,0,0,0,0,0,0,0,0,0,0,0,0,0,0,0,0,0,0,2,0,0,0,0,0,0,0,0,0,0,0,0,
0,0,0,0,0,0,0,0,0,0,1,0,0,0,1,0,1,0,0,1,0,0,0,0,0,1,2,0,1,1,2,0,
0,0,0,0,0,0,0,0,0,0,1,0,0,0,2,0,1,0,0,1,0,0,0,0,0,0,3,1,2,0,2,0,
0,0,0,0,0,0,0,0,0,0,0,0,0,0,0,0,0,0,0,0,0,0,0,0,0,0,0,0,0,0,0,0,
0,0,0,0,0,0,0,0,0,0,0,0,0,0,0,0,0,0,0,0,0,0,0,0,0,0,0,0,0,0,0,0,
0,0,0,0,0,0,0,0,0,0,0,0,0,0,0,0,0,0,0,0,0,0,0,0,0,0,0,0,0,0,0,0,
0,0,0,0,0,0,0,0,0,1,0,0,0,0,0,0,0,0,0,0,0,0,0,0,0,0,0,0,0,1,1,0),.Dim=c(17L,32L))
# observed gamete releases in the experiment. Same matrix structure as others
Gcumul<-
structure(c(0,0,1,0,0,0,0,0,0,0,0,0,0,0,0,0,0,0,0,0,0,0,0,0,0,0,0,0,0,0,0,0,
0,0,0,0,0,0,0,0,0,0,0,0,0,0,0,0,0,0,0,0,0,0,0,0,0,0,0,0,0,0,0,0,
0,0,0,0,0,0,0,0,0,2,0,0,0,0,0,0,0,0,0,1,0,0,0,0,0,0,3,0,0,0,1,0,
0,0,0,0,0,0,0,0,0,0,0,0,0,0,0,0,0,0,0,0,0,0,0,0,0,0,0,0,0,0,0,0,
0,0,0,0,0,0,0,1,0,0,0,0,0,0,0,0,0,0,0,0,0,0,0,0,0,0,0,0,0,0,0,0,
0,0,0,0,0,0,0,0,0,0,0,0,0,0,0,0,0,0,0,0,0,0,0,0,0,0,0,0,0,0,0,0,
0,0,0,0,0,0,0,0,0,0,0,0,0,0,0,0,0,0,0,0,0,0,0,0,0,0,0,0,0,0,0,0,
0,0,0,0,0,0,0,0,0,0,0,0,0,0,0,0,0,0,0,0,0,0,0,0,0,0,0,0,0,0,0,0,
0,0,0,0,0,0,0,0,0,0,0,0,0,0,0,0,0,0,0,0,0,0,0,0,0,0,0,0,0,0,0,0,
0,0,0,0,0,0,0,0,0,0,0,0,0,0,0,0,0,0,0,0,0,0,1,0,0,2,0,0,0,0,0,0,
0,0,0,0,0,0,0,0,0,0,0,0,0,0,0,0,0,0,0,0,0,0,0,0,0,0,0,0,0,0,0,0,
0,0,0,0,0,0,0,0,0,0,1,0,0,0,0,0,0,0,0,1,0,0,0,0,0,0,0,0,0,0,1,0,
0,0,0,0,0,0,0,0,0,0,0,0,0,0,2,0,1,0,0,0,0,0,0,0,0,0,0,0,2,0,1,0,
0,0,0,0,0,0,0,0,0,0,0,0,0,0,0,0,0,0,0,0,0,0,0,0,0,0,0,0,0,0,0,0,
0,0,0,0,0,0,0,0,0,0,0,0,0,0,0,0,0,0,0,0,0,0,0,0,0,0,0,0,0,0,0,0,
0,0,0,0,0,0,0,0,0,0,0,0,0,0,0,0,0,0,0,0,0,0,0,0,0,0,0,0,0,0,0,0,
0,0,0,0,0,0,0,0,0,0,0,0,0,0,0,0,0,0,0,0,0,0,0,0,0,0,0,0,0,1,0,0),.Dim=c(17L,32L))
# male number
I<-17
#female number
J<-32
# number of observed mating occasions
Kobs<-22
# maximum expected number of mating occasions
Kmax<-150
~~~

### Supplementary material Appendix 2

**Table A2.**
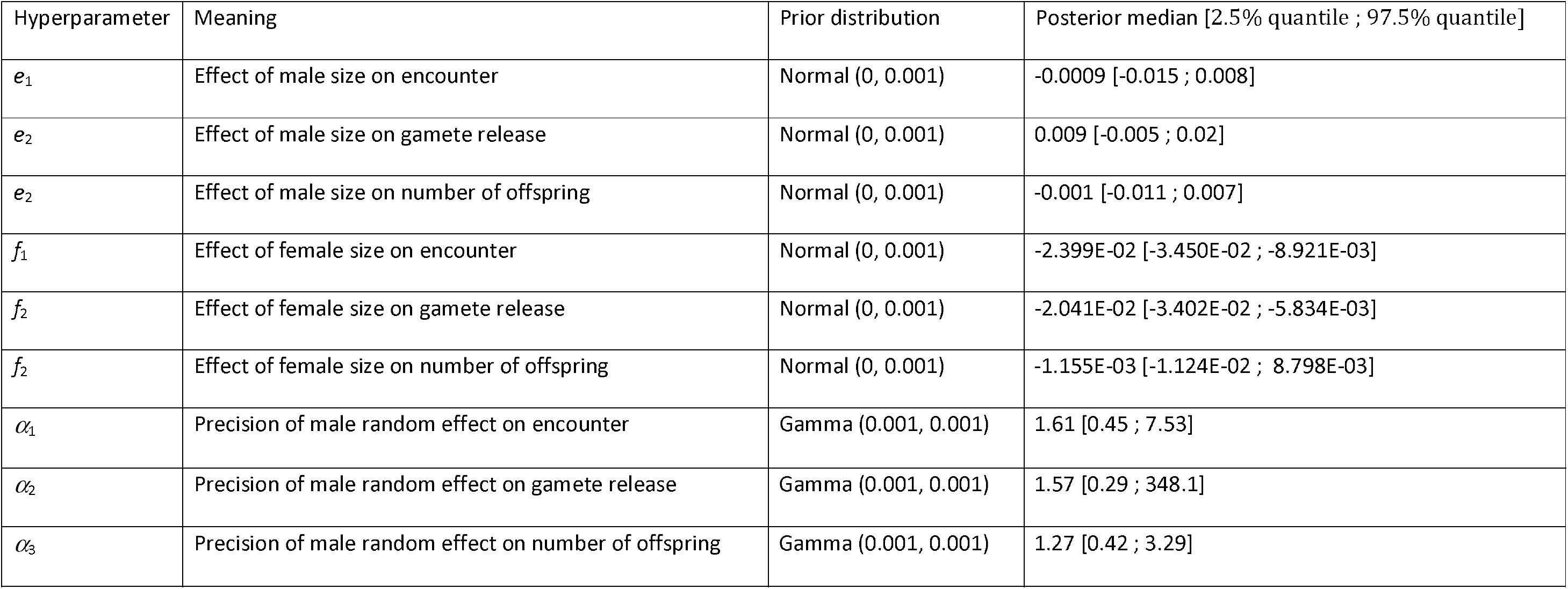
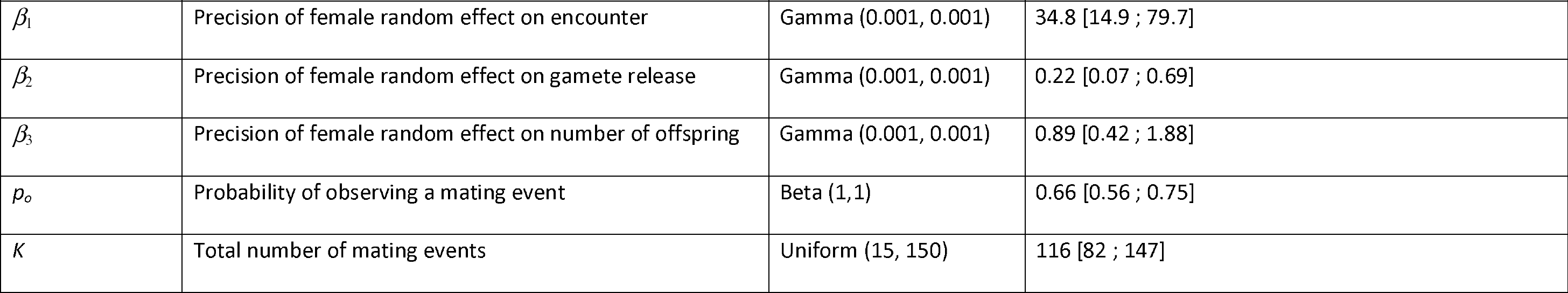
Posterior distributions for the hyperparameters of the model inferring the effect of brown trout body size on consecutive processes of roductive success.

Gauthey, Z. et al. 2017. With our powers combined: integrating behavioral and genetic data to estimate mating success and sexual selection. – Oikos 000: 000-000

